# Direct Cryo-ET observation of platelet deformation induced by SARS-CoV-2 Spike protein

**DOI:** 10.1101/2022.11.22.517574

**Authors:** Christopher Cyrus Kuhn, Nirakar Basnet, Satish Bodakuntla, Pelayo Alvarez-Brecht, Scott Nichols, Antonio Martinez-Sanchez, Lorenzo Agostini, Young-Min Soh, Junichi Takagi, Christian Biertümpfel, Naoko Mizuno

## Abstract

SARS-CoV-2 is a novel coronavirus responsible for the COVID-19 pandemic. Its high pathogenicity is due to SARS-CoV-2 spike protein (S protein) contacting host-cell receptors. A critical hallmark of COVID-19 is the occurrence of coagulopathies. Here, we report the direct observation of the interactions between S protein and platelets. Live imaging showed that the S protein triggers platelets to deform dynamically, in some cases, leading to their irreversible activation. Strikingly, cellular cryo-electron tomography revealed dense decorations of S protein on the platelet surface, inducing filopodia formation. Hypothesizing that S protein binds to filopodia-inducing integrin receptors, we tested the binding to RGD motif-recognizing platelet integrins and found that S protein recognizes integrin α_v_β_3_. Our results infer that the stochastic activation of platelets is due to weak interactions of S protein with integrin, which can attribute to the pathogenesis of COVID-19 and the occurrence of rare but severe coagulopathies.

## Introduction

In 2019 a novel member of the *Coronaviridae* family was identified to cause a respiratory illness associated with an outbreak to a global extent^1,2^. The severe acute respiratory syndrome coronavirus 2 (SARS-CoV-2) likely emerged from a zoonotic transmission similar to previous epidemic pathogens SARS-CoV and MERS-CoV, and is the origin of the Corona virus disease 2019 (COVID-19) pandemic^3^. SARS-CoV-2 is an enveloped, positive-sense single-stranded RNA virus and belongs to the genus of betacoronaviruses. It is closely related to the bat coronavirus RatG and shows 79% homology to its predecessor SARS-CoV^4,5^. Common symptoms are cough, pneumonia, and dyspnea, usually associated with a mild or moderate infection^6,7^. A severe course of the disease can have dramatic outcomes like cardiovascular complications, respiratory failure, systemic shock, and multiple organ failure, leading to life threatening conditions and potential death^8,9^.

COVID-19 is associated with abnormalities in blood coagulation in severe cases. SARS-CoV-2 is detected in the blood samples collected from COVID-19 patients^10^. Although detected viral load is generally low, the amount of virus present in the plasma correlates with the severity of COVID-19^11,12^. In a study observing the clinical aspects of COVID-19, 59.6% of the COVID-19 patients had viral loads in their blood. Particularly, in critical patients, a constant high amount of viral load (176 copies/ml) was observed, in contrast to the patients with mild cases (81.7 copies/ml)^13^. A low count of platelets (thrombocytopenia) together with the development of disseminated intravascular coagulation, myocardial infarction and non-vessel thrombotic complications are commonly observed in COVID-19 patients^8,14,15^. Platelets isolated from COVID-19 patients have also shown abnormalities such as hyperactivity and an increase in their spreading behavior^16^. The causes of these abnormalities have been hypothesized as cytokines, antiphospholipid antibodies, interactions with other immune cells, and direct interaction between SARS-CoV-2 and platelets^17–22^. Furthermore, isolated platelets from healthy donors mixed with SARS-CoV-2 or the SARS-CoV-2 spike (S) protein show a faster thrombin-dependent clot retraction and activate platelets independent of thrombin with upregulation of signaling factors^21^. A recent study further suggests the involvement of thrombin and tissue factor (TF) in the hyperactivation of platelets^23^.

Owing to its relevance for the pathogenesis of SARS-CoV-2, S protein is central to the understanding of the molecular mechanisms of the SARS-CoV-2 infection. The petal shaped S protein forms a trimer that protrudes out of the viral membrane surface^24,25^, and it is poised to engage with host cell receptors. SARS-CoV-2 and SARS-CoV S proteins show an overall similarity of 76%, although they have specific differences that impact their functions such as a furin cleavage site (PRRAR) unique to SARS-CoV-2, leading to its increased pathogenicity^26–28^. Since the beginning of the pandemic, continuous mutations have been accumulated in S protein, making it challenging to battle infections and contain disease outbreaks.

The SARS-CoV-2 S protein consists of a S1 (residues 14-685) and a S2 (residues 686-1273) subunit, which are separated by host cell proteases^29^. The receptor binding domain (RBD) in the S1 subunit is responsible for the attachment to host cells. Earlier structural studies by cryo-electron microscopy (cryo-EM) revealed open and closed states of S protein in the trimer^25,30^. In the open state, in which RBDs are uplifted thereby revealing the receptor binding motif (RBM), S protein captures its major receptor angiotensin-converting enzyme 2 (ACE2)^25,30^. In addition to ACE2, S protein is suggested to interact with several other host receptors including neuropilin-1 (NRP1)^31^ and CD147^32,33^. Interestingly, SARS-CoV-2 is the only betacoronavirus containing an RGD (Arg-Gly-Asp) tripeptide motif in the RBD, which is typically recognized by several members of the integrin membrane receptor family^34,35^. Bat-SL-CoVZC45 contains a RGD motif within the S protein but not within the RBD^36^. Initial studies observed the involvement of integrin in the SARS-CoV-2 entry^37,38^ and the binding of integrins to S protein^39,40^. However, these findings await further validation and in-depth analysis. While there is fragmented information from ultrastructural, pathological, and patient studies connecting the SARS-CoV-2 severity to the impact of its spike protein on platelets, very little is known what causes the coagulation of platelets in the presence of SARS-CoV-2. Moreover, it is still under debate if ACE2 is present on the platelet surface, and therefore, it is not clear what the direct effect of SARS-CoV-2 on platelets is^16,19,21^.

In this study, we probed the direct interaction of the SARS-CoV-2 S protein with platelets and visualized its effect on platelet morphology using live imaging and cryo-electron tomography (cryo-ET). In the presence of S protein, extensive elongation and increased spreading of platelets were observed. Notably, a population of abnormally shaped platelets resembling a proplatelet-like appearance was found, indicating an impact on the cytoskeleton-dependent platelet maturation process^41^. Cryo-ET observations revealed actin-rich filopodia formation at the end of the elongated platelet and unexpectedly, there is a dense decoration of S protein on the filopodia surface. An orientation analysis revealed that S protein binds to the membrane surface with various angular distributions. Furthermore, based on the correlation of S protein binding and the filopodia formation, the interaction of platelet surface receptors and S protein was assessed *in vitro*. We found a weak but direct interaction of platelet residing RGD ligand integrin receptors α_v_β_3_ and α_5_β_1_ with S protein but not with integrin α_IIb_β_3_. Our results shed light on the abnormal behavior of platelets leading to coagulopathic events and micro-thrombosis caused by SARS-CoV-2 infection.

## Results

### Platelet deformation in the presence of SARS-CoV-2 S protein

One of the major pathological symptoms in COVID-19 patients is an abnormal platelet behavior. COVID-19 patients exhibit conditions such as thrombocytopenia, microvascular thrombosis, and coagulation, leading to the hypothesis that SARS-CoV-2 may directly cause platelet malfunctions^8,14,15^. To assess the direct effect of SARS-CoV-2 S protein, we isolated platelets from healthy de-identified human blood donors and tested their morphological changes in the presence of S protein. Under several extracellular matrix proteins (ECMs; i.e. fibronectin and collagen I) or poly-L-lysine, platelets occasionally adhered to ECMs without S protein (Fig. 1, Movie S1). Discoid-shaped platelets were predominant in these control conditions without S protein (Fig. 1A-C; “control”). However, in the presence of S protein, we observed the deformation of platelets to elongated morphologies (Fig. 1A-C, panel “spike”). The deformation of platelets was quantified by the axial ratio (ratio of the lengths of the longer to shorter axis, Fig. 1D), showing a median value of 1.960, 1.565 and 1.786 with collagen I, poly-L-lysine and fibronectin respectively, while 1.687, 1.491 and 1.756 under control conditions without addition of S protein. The corresponding circularities in the presence of S protein decreased to 0.559, 0.649 and 0.613 with collagen I, poly-L-lysine and fibronectin respectively, compared to 0.686, 0.668 and 0.650 under control conditions (Fig. 1E). S protein-induced elongation of platelets was observed under all coating backgrounds, indicating that the morphology change of platelets is a direct effect of S protein. The deformation of platelets reached to an extent of 32 μm in the long axis and 1.4 μm in the short axis in extreme cases (Fig. S1), though precise boundaries of platelets cannot be measured due to the limitations of the light microscopic resolution. The extreme elongation of platelets (Fig. S1) indicates the presence of proplatelets, a platelet precursor present in the circulatory system^42,43^. Typically, proplatelets are processed by cytoskeletal remodeling and abscise into smaller platelets within the circulatory system^42,43^. Our observation suggests that S protein may influence the maturation of platelets by acting on the cytoskeleton-based process necessary for proplatelet division. Occasionally, we observed a proplatelet shape, (Fig. S1, white arrowhead), resulting in the formation of a wide hollow, ring with one or more bulges like gemstones on a ring. Interestingly, the effect of platelet deformation is less prominent in the poly-L-lysine background (Fig. 1B). We also observed less effect in the activation of platelets with S protein (Fig. 1F, Fig. 2D) with poly-L-lysine. As poly-L-lysine is not a platelet-specific adherence factor, it suggests that the deformation of platelets may be further aided by the binding to extracellular agents. To test the effects of S protein in the activation of platelet, a solid-phase sandwich ELISA assay and western blotting were performed. In the ELISA assay, platelet factor 4 (PF4) was measured to test the secretion of alpha-granules, a marker for the activation of platelets^44^. The result showed an increase in secreted PF4 in platelet samples in the presence of S protein (Fig. 1G). Western blot analysis showed an increase in phosphorylated focal adhesion kinase (pFAK) as well, which responds to the signaling cascade of focal adhesion pathway^45^. Interestingly, while pFAK was detected from platelets that adhered to the plate surface (1.45 ± 0.51 times higher than control without S protein), floating platelets did not show a detectable level of pFAK (Fig. S1B-D). This indicates that the deformation of platelets is a reversible process that has little or only local influence.

**Figure 1.**
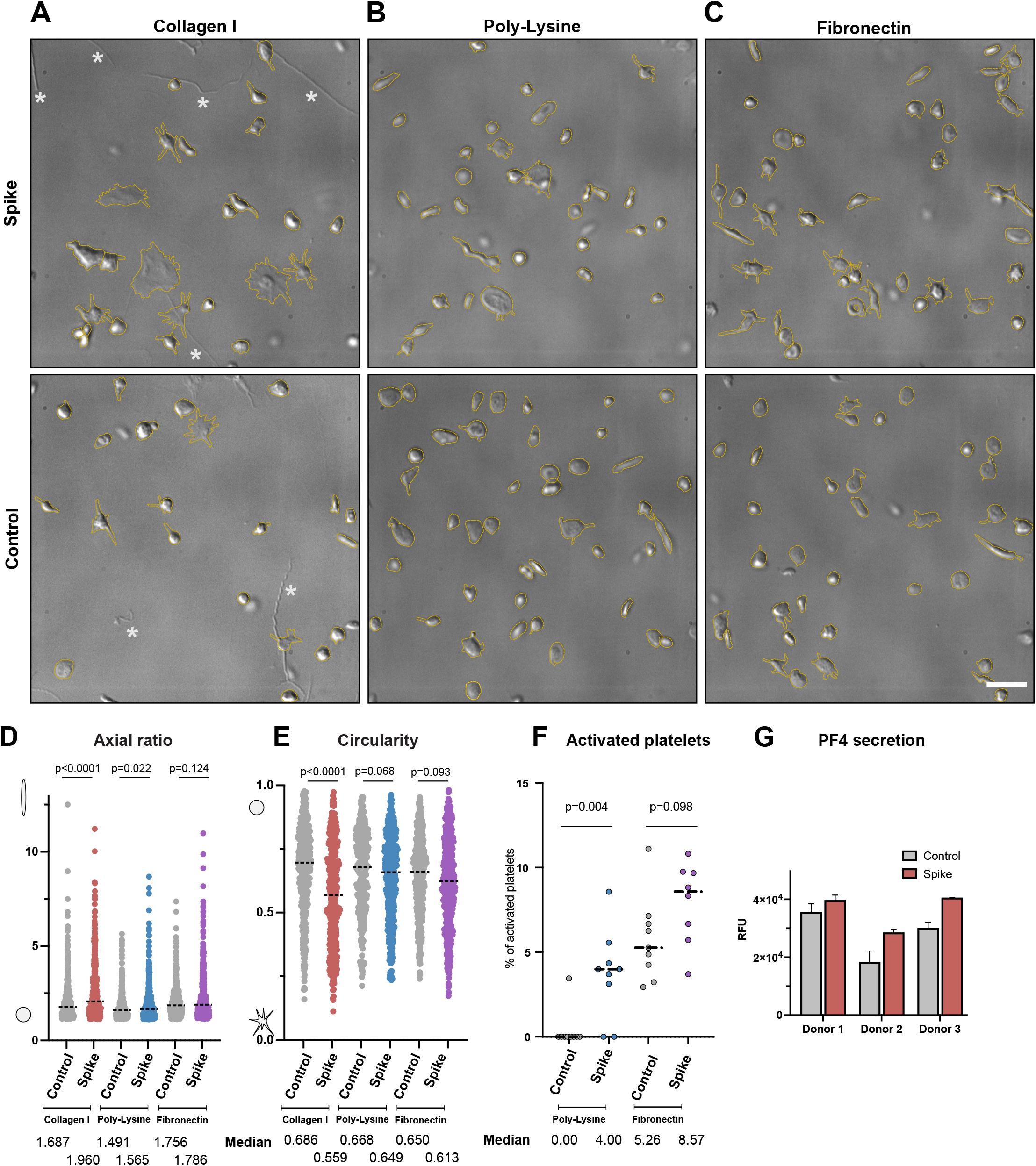
Comparison of platelet morphology with and without SARS-CoV-2 S protein. (A) DIC images of platelets without (Control) and pre-incubated with 20 μg/ml S protein (Spike) on a collagen I support. The platelet shape is outlined in dashed yellow. * indicates collagen I fibers. (B) DIC images of platelets without and pre-incubated with S on a poly-L-lysine support. (C) DIC images of platelets without and pre-incubated with S protein on a fibronectin support. Scale bar: 5 μm (A-C). (D) Quantification of the axial ratio of platelets (major axis/minor axis) on different coated surfaces, without and in the presence of S protein The median axial ratio is shown below the corresponding violin plot. The significance was determined by Mann-Whitney U test. (E) Quantification of the circularity of platelets on different coated surfaces, without and in the presence of S protein. The median circularity is shown below the corresponding violin plot. The significance was determined by Mann-Whitney U test. (F) Comparison of platelet activation, incubated with and without S protein, on Poly-L-Lysine (left), and on Fibronectin (right) Platelets with amoeba-like morphologies were defined as activated platelets. The significance was determined by Mann-Whitney U test. (G) Sandwich ELISA assay detecting PF4 release in the absence and presence of S protein.

**Figure 2.**
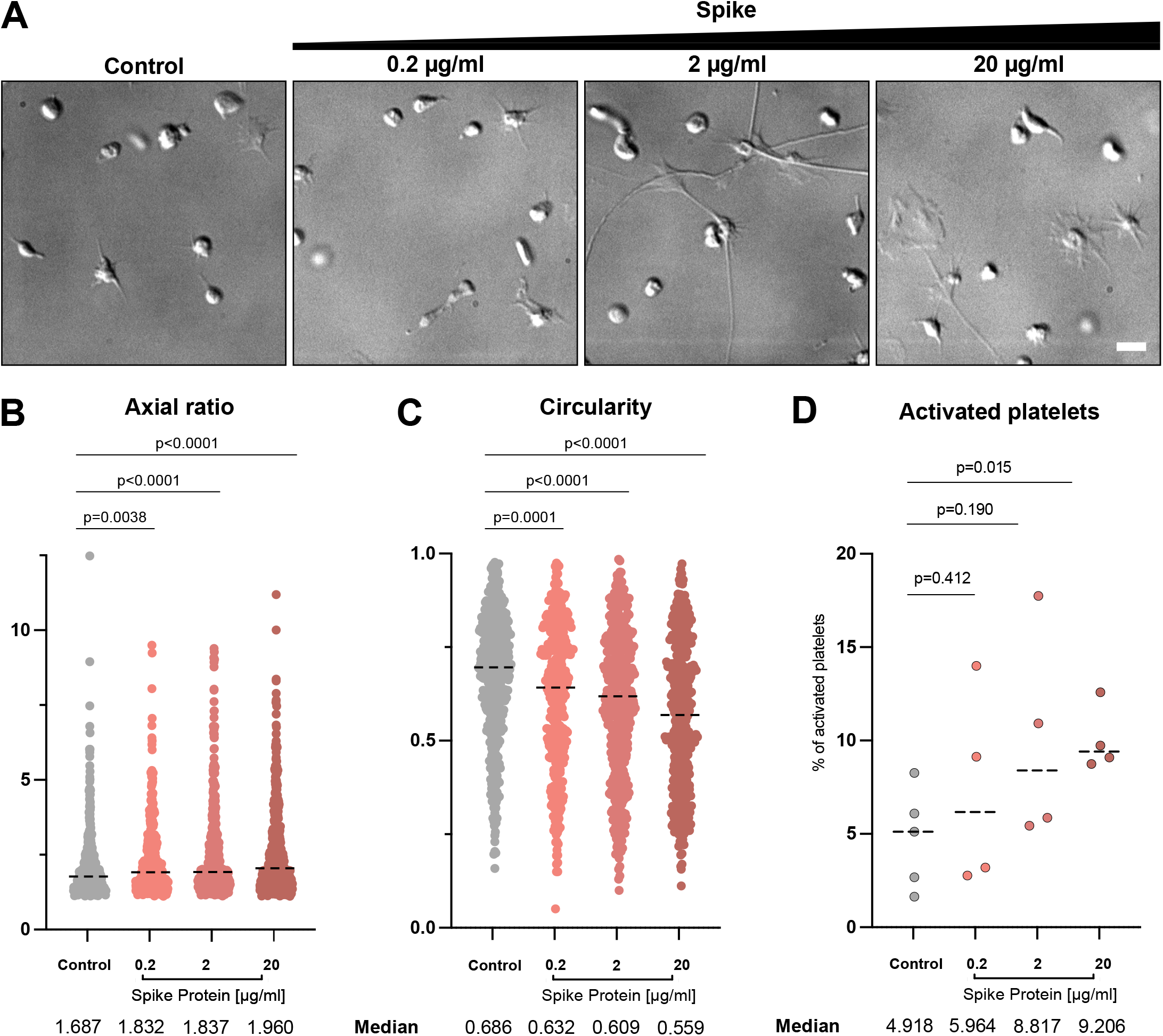
Platelet morphology depending on SARS-CoV-2 S protein concentration. (A) DIC images of platelets in the presence of different amounts of S protein plated onto collagen I-coated surfaces. Scale bar: 5 μm. (B) Quantification of the axial ratio of platelets on collagen I-coated surfaces, without and in the presence of different S protein concentrations. The median axial ratio is shown below the corresponding violin plot. The significance was determined by Mann-Whitney U test. (C) Quantification of the circularity of platelets on collagen I-coated surfaces, without and in the presence of different S protein concentrations. The median circularity is shown below the corresponding violin plot. The significance was determined by Mann-Whitney U test. The plots for control and in the presence of 20 μg/ml spike protein in (B) and (C) are same as those in Figure 1D and E, respectively. (D) Quantification of platelet activation on Collagen I depending on S protein concentration. The significance was determined by Mann-Whitney U test.

On the surface of SARS-CoV-2, S protein is arranged with an average distance of 35 nm to its neighboring S protein with an estimated number of 24 S proteins per virus particle, though the reported distribution has no apparent geometrical order^46^. In this way, S proteins are effectively locally concentrated on the viral surface. To reflect the local concentration effects and test the influence of the concentration of S protein on the deformation of platelets, various concentrations of S protein were added to platelets in tenfold steps from 0.2 μg/ml to 20 μg/ml (0.47 nM to 47 nM) and the morphology changes of platelets were assessed. We observed an increase in the elongation as well as activation of platelets in accordance with the concentration of S protein (Fig. 2). While S protein was able to cause the deformation of the platelets even at the lowest concentration (0.2 μg/ml, or 0.47 nM), more dominant effects were observed under high concentrations. Our observation suggests that higher concentrations of S protein have a more pronounced impact on platelets, which could mimic locally concentrated S protein on the viral surface.

### Cryo-ET analysis shows S protein densely decorated on the platelet surface

To gain molecular insights into the morphological changes of platelets in the presence of S protein, we performed a cryo-ET analysis of platelets under a collagen type I background, both in the presence and absence of S protein (Movie S2). Platelets incubated with S protein revealed extensive deformations (Fig. 3A-3C, highlighted in purple, Fig. S2) consistent with the light microscopic observations, while intact platelets exhibit their typical disc-like morphology (Fig. 3D–3F, highlighted in yellow). High magnification observation (33000x) revealed that the elongated morphology of platelets was facilitated by the remodeling of actin, forming filopodia-like architectures as narrow as 43 nm (Fig. S2). Within the filopodia that we analyzed (Fig. 3G and 3I), we found actin filaments assembling into tightly packed bundles, comparable to those seen in pseudopodia of untreated platelets^47^. Accompanied by longer actin filaments running parallel to the axis of the filopodia, shorter ones were bridging between the longer filaments and the membrane surface (Fig. 3H–3J). The angular distribution of actin segments shows two peaks, namely at 10 and 80 degrees (Fig. 3H, arrows), those of 10° corresponding to the actin filaments that are running along the filopodia axis, while those of 80° connecting between the actin bundles and membranes. The difference of 70° between peaks reinforces the notion that the connections are made by the Arp2/3 complex, which is known to mediate actin branching at 70°^48^. Control platelets showed filopodia-formation exclusively when contacting collagen I fibrils (Fig. 3D–3F). Interestingly, in the presence of S protein, the morphological changes of platelets and their filopodia-like formation did not require the attachment to collagen I (Fig. 3A–3C). This indicates that S protein itself has a contribution to the surface activation of platelets. Unexpectedly, we identified dense S protein densities decorating the external membrane surface of the filopodia protrusions, indicative of S protein (Fig. 3K, “+”, Movie S2). In contrast, we only observed shorter faint densities of endogenous membrane receptors on the surface of intact platelets (Fig. 3K, “-”).

**Figure 3.**
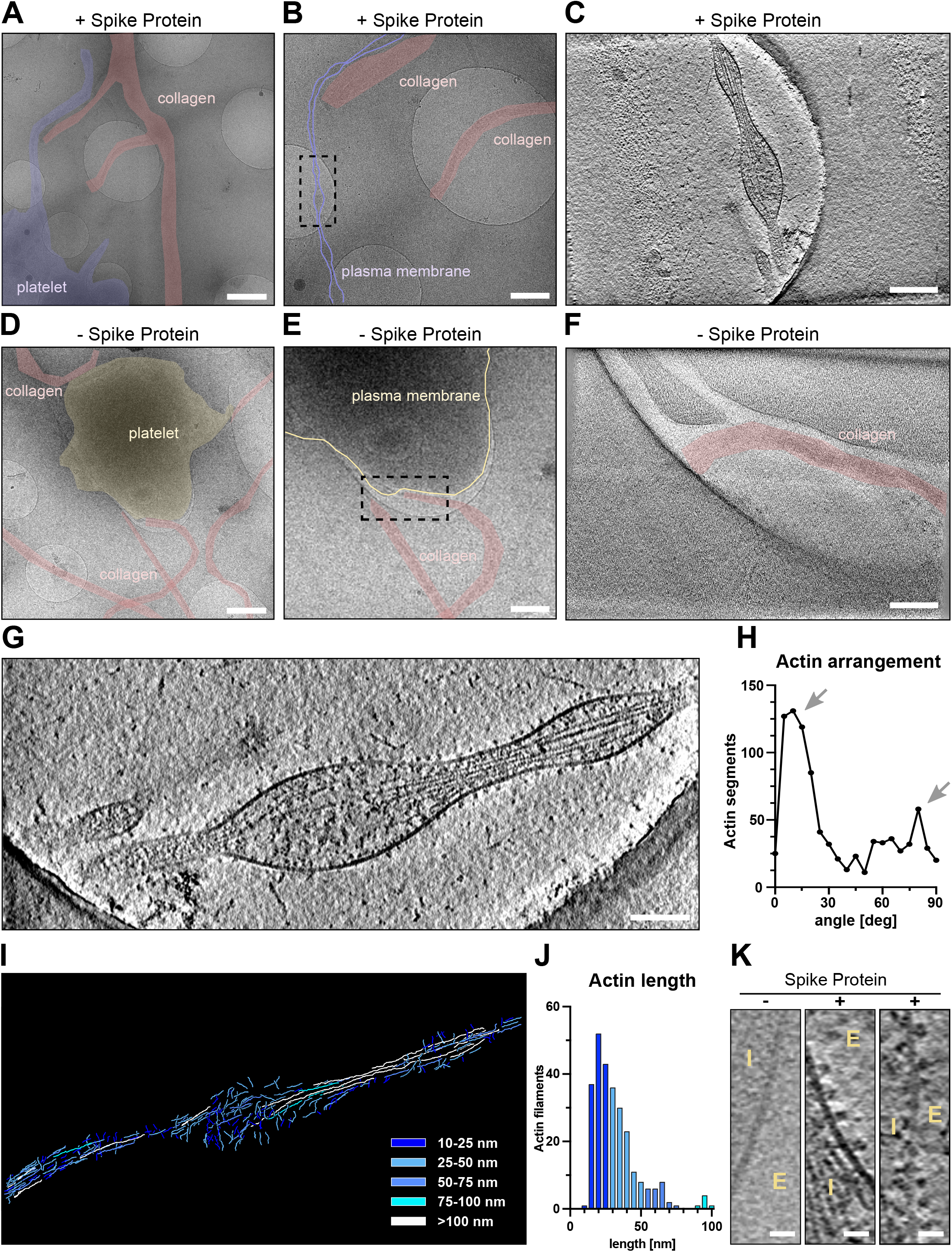
Cryo-electron tomograms of platelets alone and in the presence of SARS-CoV-2 S protein on a collagen I support. (A) and (B) Low magnification views of a platelet in the presence of S protein on collagen I. The dashed box in B represents the area of tomographic data collection in C. (C) Tomographic slice of the platelet protrusion in the presence of S protein. The platelet is indicated in purple, collagen I fibers in red. (D) and (E) Low magnification views of a platelet on collagen I. The dashed box in E represents the area of tomographic data collection in F. (F) Representative slice of the reconstructed tomogram of platelet plasma membrane. The platelet is indicated in yellow, collagen I fibers in red. (G) Magnification of the filopodial structure from C with actin filaments running along the protrusion. (H) Angular arrangement of actin filaments along the platelet protrusion. (I) Traced actin filaments of the tomographic reconstruction in G. Actin filaments are color-coded by length of blue to white. (J) Length distribution of traced actin filaments depicted in I. Actin filaments ≥ 100 nm (12 in total) are not represented in the graph. (K) Magnified views on the platelet plasma membrane without and in the presence of S protein (E - extracellular, I–intracellular). Scale bars: (A),(D) = 1 μm; (B),(E) = 0.5 μm; (C),(F),(G) = 200 nm; (K) = 20nm.

### SARS-CoV-2 S protein binds to the platelet membrane surface flexibly

To further analyze the S protein densities on the platelet membrane surface, we manually selected and extracted the densities from 8 tomograms and analyzed them using subtomogram averaging approaches (Fig. 4A). To facilitate a focused alignment of the decorating protein without the influence of the membrane density, the membrane signal was subtracted using PySeg^49^. We obtained a 3D-averaged density at a resolution of 13.8 Å (Fig. 4A, Fig. S3) showing a characteristic trimeric shape of 15 nm in size. The obtained structure agreed well with our near-atomic resolution structure of S protein using the same protein batch for single-particle analysis (Fig. S3) as well as previously published structures^25,30,46,50^, (Fig. 4A, fitted PDB 6vxx), validating the identity of S protein decorations. Furthermore, the analysis of the particles without application of C3 symmetry showed an asymmetric uplift of the tip of the S1 surface that connects to the extra density (Fig 4A, right). This shows that one of the three RBD domains of S protein is lifted up upon its binding to the host cell receptors. The extra density connected to S protein represents the density from the host cell, likely the platelet surface receptor recognized by S protein (Fig. 4I).

**Figure 4.**
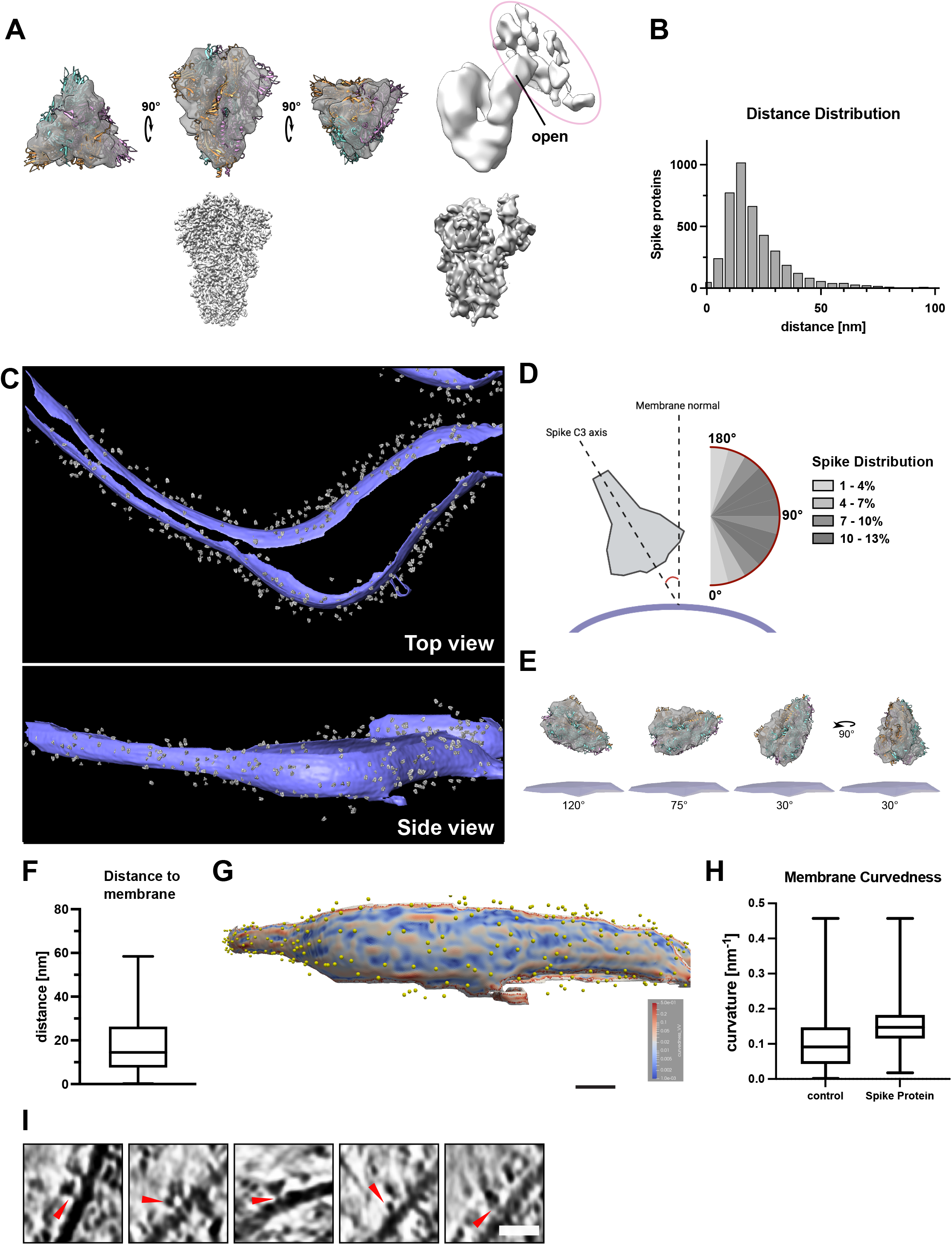
SARS-CoV-2 S protein reconstruction and membrane decoration analysis. (A) Top-Left: Structure of S protein with closed conformation fitted in the subtomogram reconstruction. Top-Right: Structure of S protein calculated without C3 symmetry, revealing the uplifted RBD domain connected to additional densities from the host platelets. The additional densities connected to the open RBD domain is circled in magenta. Bottom-left: SPA-based structure of S protein in the closed conformation. Bottom-right: SPA-based structure of S protein in the open conformation. (B) Nearest neighbor distance distribution of S protein densities on the platelet surface membrane. The distances are calculated using the originally manually picked coordinates. The median distance between two S protein is 27.3 nm (C) Densities of the reconstructed S protein back-plotted to the segmented platelet plasma membrane (purple - platelet plasma membrane, gray - S protein). The tomogram lacks top and bottom due to the “missing wedge” effect of tomographic data collection. (D) Orientation of S protein (grey) on the membrane surface (purple). The scheme depicts the angle determination of S protein C3 axis and the normal of the platelet plasma membrane. The range from 45-120° was observed to be favorable for S protein interaction with the platelet surface. (E) Schematic depiction of S protein orientation in different angles towards platelet plasma membrane. (F) Distance of S protein from the membrane. The median distance from the center of S protein to the membrane is 16 nm. The box plot represents 25 and 75 percentiles (8.6 and 27 nm, whiskers). (G) Visualization of the platelet plasma membrane curvedness. The yellow spheres indicate the position of S protein on the platelet surface. The top and bottom edge of the segmented membrane was excluded from the estimation and is colored in grey. (H) Membrane Curvature comparison of S protein bound and surface protein free areas on the platelet plasma membrane. The box plots represent 25, median and 75 percentiles. Control: 25% 0.043, median 0.091 and 75% 0.15. + S protein: 25% 0.11, median 0.14 and 75% 0.17. (I) Additional densities (red allow-heads) between picked S protein and platelet plasma membrane. Scale bar: (I): 20nm.

To assess how S protein recognizes the platelet surface, the alignment parameters of the individually analyzed S protein densities were applied to the 3D average and plotted back to the original tomograms (Fig. 4B-H). The distribution analysis of neighboring S protein showed that the peak population had a distance of 27.3 nm (median) apart, but some molecules were also more sparsely distributed (Fig. 4B). This measurement corresponds to a density of one S protein on a surface area of up to 585 nm^2^ (a radial surface of 585 nm^2^ is covered by one S protein), although no apparent periodical distribution was detected. Judging from the diameter of S protein (~17 nm), neighboring S protein closely located next to each other. S protein bound to the platelet membrane at a distance of 16 nm between the center of S protein to the membrane surface (Fig. 4F) and interestingly, with a wide range of angular distribution (Fig. 4D-E) with respect to the membrane surface, indicating its flexible attachment to the platelet surface. In addition, S protein binds to a slightly more curved membrane surface (Fig. 4G and 4H). This may be reflected by the fact that the binding of S protein induces filopodia formation with a membrane protrusion. Taken together, these results indicate that S protein approaches the platelet surface from various geometrical orientations to accommodate and enhance the docking to the membrane surface. Similarly, a broad angular distribution of S protein has been observed from the viral surface, due to several kinked points in the stalk region^46,50^. Together with our observation, it suggests the orientational adjustments from both sides of S protein, namely the receptor binding S1 subunit and the stem side at the root on the virus, maximize the efficiency of the attachment of S protein to the host cell receptors. Consistent with the observed additional density on the lifted RBD domain (Fig. 4A, right), some of the tomograms showed extra densities bridging between plasma membrane and S protein (Fig. 4I, red arrows), presumably those of platelet receptors recognizing S protein. However, subtomogram analysis only yielded a faint density (Fig. 4A, right) without features, also suggesting a flexible attachment of S protein to its receptor on the membrane surface.

### Platelet deformation in the presence of pseudotyped viral particles

After characterizing the effects of S protein on platelets, we hypothesized that locally concentrated S protein on a globular viral surface would be advantageous for increasing the local concentration and evaluated the influence of SARS-CoV-2 pseudo virus-like-particles (VLPs) on platelet deformation. We either generated or obtained SARS-CoV-2 pseudotyped VLPs that are fully intact vesicle-like entities as validated by negative staining EM (Fig. S4A) and by their ability to infect HEK-293T-hACE2 cells (HEK-293T cells constitutively expressing ACE2 receptor, Fig. S4B-S4C). The viral titer determined by flow cytometry was approximately 10^4^-10^6^ particles/ml, comparable to the reported preparation of SARS-CoV-2 pseudo VLPs^51^, however, it was low compared to VSV-G based lentiviruses. This low titer did not allow us to readily detect changes in platelet morphologies by live platelet imaging. However, we were able to find an example of a particle located in close proximity to a platelet filopodium (Fig. S5B-S5F). Cryo-electron tomography revealed that the closest distance between this particle and the membrane surface of the filopodium was 20 nm (Fig. S5G and S5H), similar to that measured for S protein alone (Fig. 4F 16 nm, from the center of S protein to membrane). The cross-section views of the particle showed decorations of proteins on the membrane surface (Fig. S5I, red arrowheads), altogether suggesting that this vesicle may be a bound VLP. In comparison, we also found examples of extracellular vesicles with a similar shape and size (Fig. S5A) that appeared to originate from intracellular vesicles, i.e. exosomes, containing alpha granules (Fig. S5A, left) or budding out from the concave surface of plasma membrane, instead of filopodia and indicative of vesicle release through fusion of lysosome and plasma membrane. These results corroborate our *in vitro* data of purified S protein inducing morphological changes in platelets.

### Integrin receptors recognize SARS-CoV-2 S protein

Several cell receptors were reported to recognize S protein. However, the presence of ACE2, the major S protein receptor, on the platelet surface is still inconclusive^16,19,21^.Therefore, the relevance of the ACE2 receptor for platelet malfunction is still an open question. In contrast, integrin receptors are the major class of receptors expressed in platelets. Considering our structural analysis (Fig. 4) and the possibility that ACE2 is not abundantly expressed on platelets, we hypothesized that S protein may directly recognize integrin receptors. Interestingly, the RBD domain of S protein contains a stretch with an “RGD” motif, which is a common motif among integrin ligands^35^ and a direct interaction of tissue integrin α_5_β_1_ and SARS-CoV-2 S protein has been shown^39^. We therefore tested the binding of S protein to known platelet integrin receptors α_IIb_β_3_, α_v_β_3_, and α_5_β_1_, enriched in the tissue but also expressed on platelets, all recognizing the RGD ligand motif. We used ELISA-like solid-phase equilibrium binding assays to detect the interaction of S protein with integrins (Fig. 5A). We detected the binding of integrin α_5_β_1_ and α_v_β_3_ to S protein, while integrin α_IIb_β_3_ does not have an apparent interaction with it (Fig. 5B). The extent of binding is most prominent with integrin α_v_β_3_, while integrin α_5_β_1_ showed only a weak interaction. However, it should be noted that the observed binding of tested integrins was much weaker (less than 10-fold) compared to those for physiological integrin ligands: vitronectin for α_v_β_3_, fibrinogen for α_IIb_β_3_ and fibronectin for α_5_β_1_. Encouraged by our results, we tested the effect of platelet activation in the presence of cilengitide, a cyclic RGD pentapeptide^52^ that blocks the binding of integrin to RGD motif-containing extracellular ligands, and indeed the activation was reduced (Fig. 5C). These observations generally agree with a recent discussion of the relevance of integrin recognition by SARS-CoV-2 for vascular dysregulation^38^.

**Figure 5.**
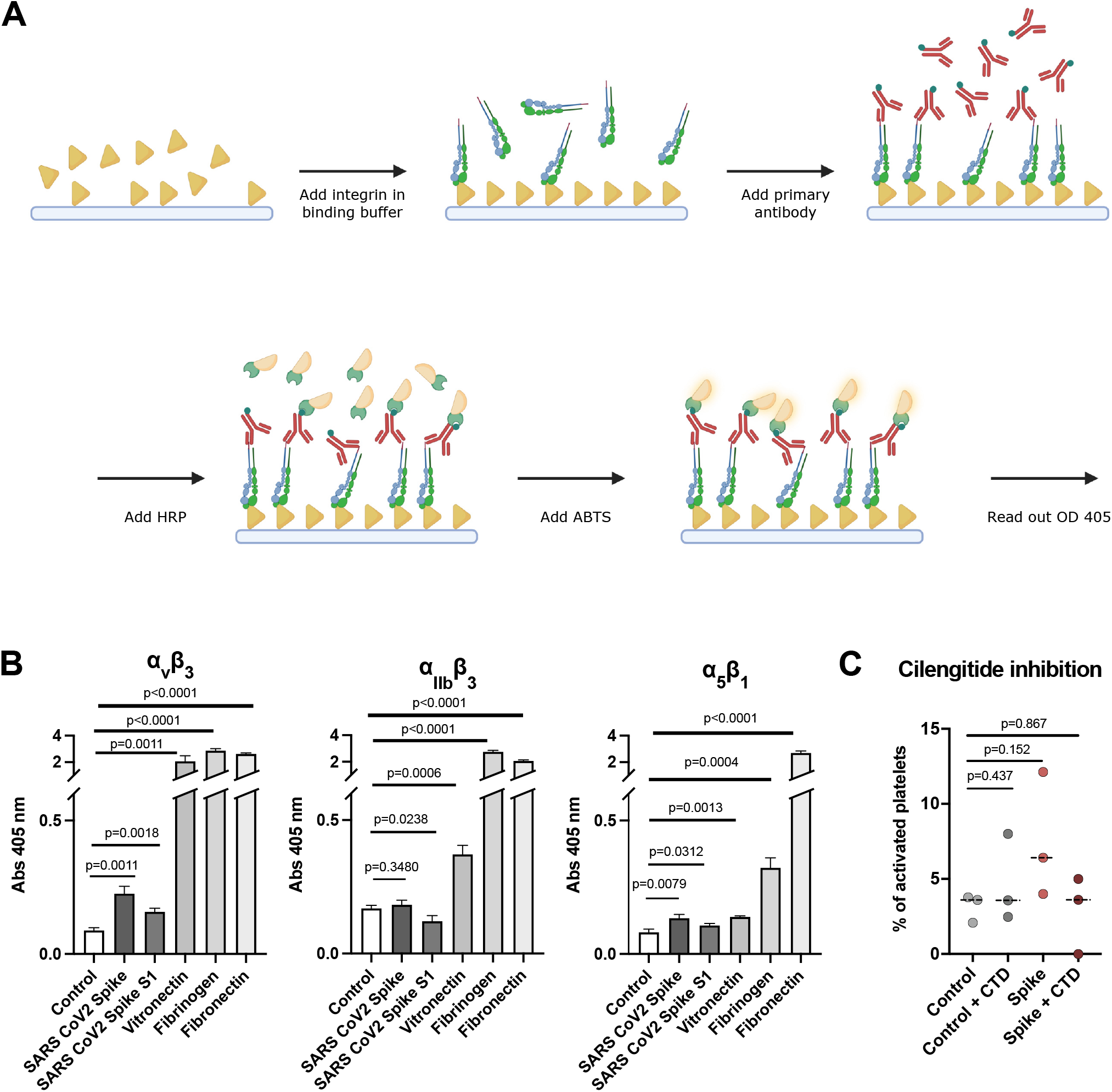
Interaction of integrin receptors with SARS-CoV-2 S protein and various ECM proteins. (A) Scheme of the experimental setup. Immunoplates were coated with either S protein or ECM proteins. The ligands were incubated with various integrin-velcro constructs. Biotinylated anti-velcro polyclonal antibody, subsequently coupled to Strepanvidin-HRP, was used to label ligand-bound integrins. Detection of the binding was measured at 405 nm, 10 min after addition of ABTS. The scheme was created with Biorender.com. (B) Binding of integrins α_v_β_3_, α_IIb_β_3_, and α_5_β_1_ to S protein and their physiological ECM ligands: α_v_β_3_ - vitronectin, α_IIb_β_3_ - fibrinogen, α_5_β_1_ - fibronectin. Data are from a representative experiment out of three independent ones, and shown as mean ± SD. The significance was determined by an unpaired t test. (C) Quantification of platelet activation on Collagen I depending on S protein concentration and integrin inhibitor cilengitide. The significance was determined by Mann-Whitney U test.

## Discussion

SARS-CoV-2 has shown unique pathological symptoms that can lead to a wide range of coagulopathic events in severe cases. In our study, we probed the direct effect of S protein to the change in morphology of platelets at a molecular level, and for the first time, we directly visualized the binding of S protein to the platelet surface (summarized in Fig. 6). We hypothesized that the binding of the SARS-CoV-2 is mediated by integrin receptors based on the following reasons; 1) the activation of platelets is governed by filopodia formation, 2) filopodia formation is initiated by integrin receptors, 3) the major receptors on the platelets are integrin receptors and 4) SARS-CoV-2 S protein contains a “RGD” sequence in the RBD, which is recognized by a subtype of integrin, and therefore we tested the interaction of platelet-expressed integrins with S protein. Our integrin inhibition experiment using cilengitide and *in vitro* solid-phase binding assays support this hypothesis, particularly with the possibility that S protein recognizes integrin α_v_β_3_. The binding of S protein to integrin was much lower compared to the interaction of integrins with their physiological ligands, and interestingly, we did not detect the binding to the major platelet integrin α_IIb_β_3_. Previously, an increased binding of the activated integrin α_IIb_β_3_ antibody PAC-1 to platelets was observed in the presence of S protein^21^. This may be due to an inside-out effect, in which the outside-in signaling is activated by the direct binding of S protein to integrin α_v_β_3_ and in turn, α_IIb_β_3_ would get activated through the intracellular signaling (inside-out). We surmise that the weak affinity of S protein to platelet integrin receptors and the reversible binding, may reflect the fact that blood clotting defects observed in patients are rare complications and occur in severe cases of COVID-19. However, here we should also note that there are other receptors on platelets that may also be accountable for the interaction with S protein^33,53^ and combinatory effects of the binding of S protein to multiple receptors may also occur.

**Figure 6.**
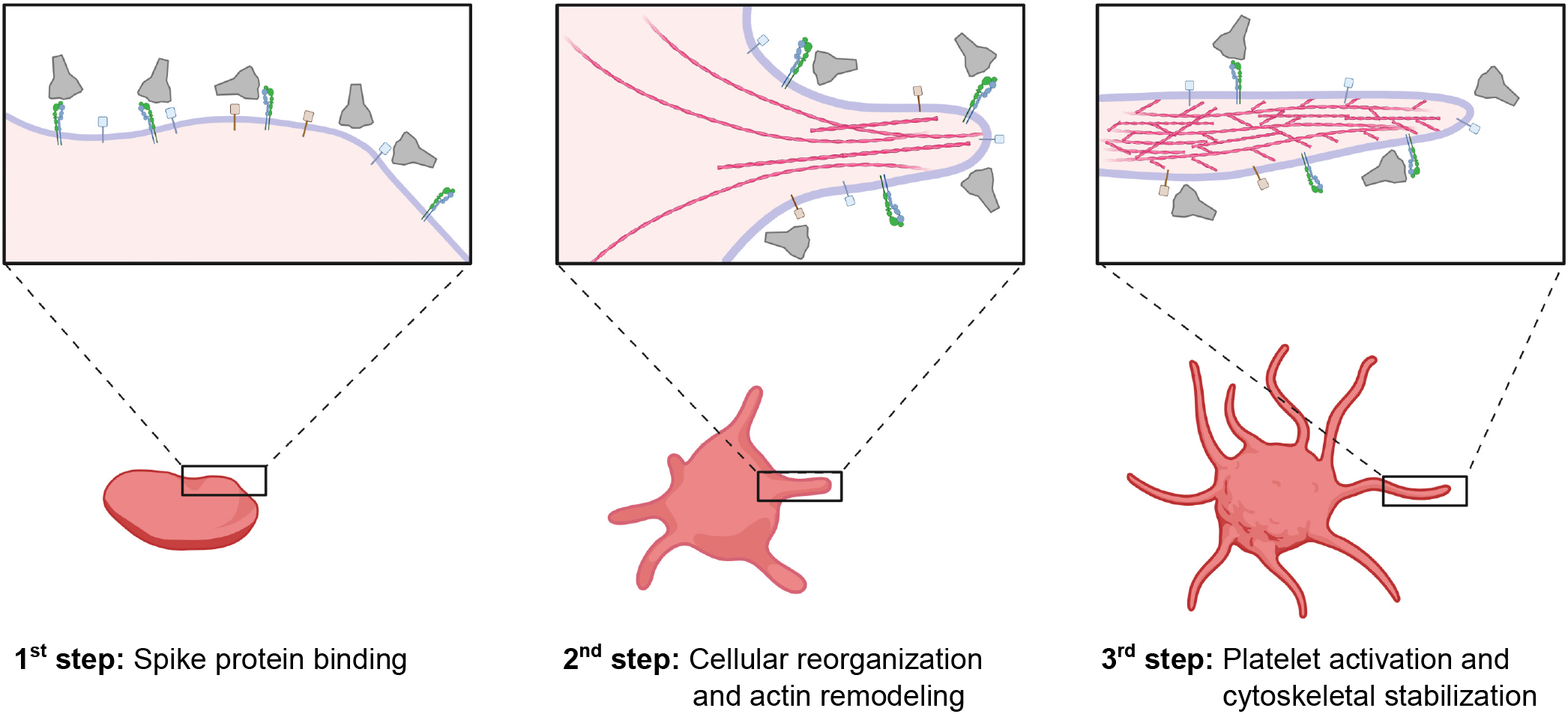
Schematic representation of potential SARS-CoV-2 S platelet interaction. First, S protein binds to receptors on the platelet surface, causing the deformation and priming the activation. Protrusions are forming as a consequence of actin remodeling. This leads to the activation of platelets by the formation of filopodia and the stabilization of the cytoskeleton network. The scheme was created with Biorender.com.

SARS-CoV-2 is found in the blood stream of COVID-19 patients^10^, and an open question is how it can lead to rare but severe coagulation defects. We showed that the deformation of platelets itself does not always alter their intracellular signaling (Fig. S1), or induces activation. It rather appears that platelets exposed to S protein are primed for the activation upon further stimuli, such as the attachment to an adhesion surface. Based on this observation, we speculate that the combination of the direct binding of S protein to platelets and other identified coagulation factors may induce a synergistic and irreversible activation of platelets, leading to coagulation. During SARS-CoV-2 infection, several other procoagulant players are active, for example the formation of neutrophil extracellular traps^18^, the release of TF^23^, elevated fibrinogen levels^54^ and dysregulated release of cytokines^55^, creating a hypercoagulative environment in the context of COVID-19.

In our study, we visualized the adaptable attachment of S protein to the platelet plasma membrane with a high degree of flexibility for the engagement to continuously curved membrane surfaces (Fig. 4D and 4E). Similarly, it has been reported that the stalk domain of S protein proximal to the viral membrane surface contains three hinges, presumably allowing the flexible motion of individual S protein on the viral surface to adapt to curved host cell surfaces^50^. This dual flexibility likely increases the probability for S protein to attach to a host cell receptor, thus, allowing an efficient action of S protein to the membrane surface.

## Data availability

Tomograms of platelets in the presence of SARS-CoV-2 S protein used in the figures were deposited to the Electron Microscopy Data Bank (EMDB) with accession codes EMD-26794 (platelet protrusion shown in Fig. 3) and EMD-26796 (platelet protrusion shown in Fig. 4). Additional tomograms used for subtomogram averaging were deposited to the Electron Microscopy Public Image Archive (EMPIAR) with accession code EMPIAR-11038. The 3D map of the single particle reconstructed S protein has been deposited with the accession code EMD-26798.

## Acknowledgements

We thank the members of Mizuno Lab for discussions and help for this research. We thank William Wan for their insightful suggestions and advice for the tomographic analysis and membrane curvature quantification. We thank Ana Pasapera for the technical advice of signaling assays. The light microscopic data was collected at the NHLBI light microscopy core facility, the flow analysis was performed at the NHLBI flow cytometry core and the cryo-ET data was collected at the MICEF at the National Institutes of Health, USA and at the National Cancer Institute’s National Cryo-EM Facility at the Frederick National Laboratory for Cancer Research under contract HSSN261200800001E. The following reagents were obtained through BEI Resources, NIAID, NIH: SARS-Related Coronavirus 2, Wuhan-Hu-1 Spike-Pseudotyped Lentivirus, Luc2/ZsGreen, NR-53818. NM acknowledges the Intramural Research Program of the National Heart Lung and Blood Institute, and the National Institute of Arthritis and Musculoskeletal and Skin Diseases of National Institutes of Health, USA, for funding.

## Methods

### Platelet Isolation

Human platelets were prepared from the blood of de-identified healthy donors according to the available protocol^21^, with minor modifications as described below. Immediately after blood was drawn from the donors, it was centrifuged for 20 min at 200 g at RT. The top half of the platelet-rich plasma (PRP) was transferred to a fresh tube and gently mixed with an equal amount of HEP buffer (14 mM NaCl, 2.7 mM KCl, 3.8 mM HEPES, 5 mM EGTA, 1 μM Prostaglandin E1, pH 7.4.). To remove remaining cells other than platelets, the PRP solution was centrifuged for 20 min at 100 g. Three fourth of the supernatant was carefully transferred to a fresh tube, and platelets were pelleted for 20 min at 800 g. The supernatant was discarded, the pellet was then washed with a solution containing 10 mM sodium citrate, 150 mM NaCl, 1 mM EDTA, 1% (w/v) dextrose, pH 7.4 and resuspended in Tyrode’s buffer (134 mM NaCl, 12 mM NaHCO_3_, 2.9 mM KCl, 0.34 mM Na_2_HPO_4_, 1 mM MgCl_2_, 10 mM HEPES, 5 mM Glucose, 3 mg/ml BSA, pH 7.4). The isolated platelets were rested for 45-60 min. The platelet concentration was estimated using a hemacytometer.

### Preparation of Coating

Poly-L-Lysine (Sigma #P2636) or ECM proteins fibronectin (R&D systems #3420-001-01) and collagen-I (Chrono-Log #385) were diluted to working concentration 0.01% for Poly-L-Lysine, 10 μg/ml for Fibronectin and 25 μg/ml for Collagen I respectively. Poly-L-Lysine and fibronectin were applied to imaging chambers (μ-Slide 8 well glass bottom, ibidi #80827-90) over night at 37 °C. Collagen I coating was applied for 30 min at 37 °C. Coated imaging chambers were washed with PBS and afterwards blocked with 1% BSA in PBS.

### Platelet and S protein Incubation

S protein (Cube Biotech #28703) and isolated platelets were mixed at final concentrations of 0.2 μg/ml, 2 μg/ml and 20 ug/ml of S protein and 1.5×10^7^ platelets/ml in Tyrode’s buffer. The samples were incubated for 4 h at 37 °C, and further diluted to 0.75×10^7^ platelets/ml. Controls were prepared by adding Cube Biotech’s S protein buffer (20 mM HEPES, 150 mM NaCl, 0.01% LMNG, pH 7.5) instead of S protein.

### Differential Inference Contrast (DIC) Microscopy and Analysis

Prior to the imaging, the coated imaging chambers were fixed inside the Live-Cell Imaging chamber and the chamber and water bath temperature were set to 37 °C. The platelet and S protein mixture was transferred to the coated imaging chambers using wide orifice tips. Platelets were settled for 15 min, and subsequently live imaging was performed for 60 min, with a frame time of 2 min, except one series of experiment recorded for 58 min. Acquired frames were analyzed using FIJI^56^. The platelet shape was manually tracked and segmented every 3 frames using ‘freehand selection’ tool over the imaging period. The overall circularity and axial ratio of the platelets were quantified using the FIJI measurement plugin with options of ‘circularity’ and ‘aspect ratio’. Circularity is calculated by 4π(area/perimeter^2^). A circularity value of 1.0 indicates a perfect circle and lower values indicate an elongated polygon. Axial ratio is defined as the ratio of the major axis and minor axis of the fitted ellipse. The quantified data was analyzed using GraphPad PRISM. The measurements were done at least 3 times using platelets from at least 3 different donors.

### Platelet activation in the presence of integrin inhibitor cilengitide

Isolated platelets were pre-incubated at 111 μg/ml cilengitide (Millipore-Sigma #ML1594) for 20 min at 37°C. Subsequently, S-protein or vehicle control was added to the platelets so that the final concentration of cilengitide becomes 100 μg/ml. The platelet activation was assessed by live imaging in the same way as the DIC assay without cilengitide.

### PF4 sandwich ELISA assay

Supernatant from platelets seeded on Collagen I either incubated with or without S-protein was collected. The PF4 concentration was determined using the Human PF4 CatchPoint^®^SimpleStep ELISA^®^Kit (Abcam #ab278096), according to the manufacturer’s protocol. Prepared samples, standards and antibody cocktail were added to appropriate wells and incubated for 1 h at RT while gently shaking. Subsequently, the solution was removed and wells were washed with 1x wash buffer PT. CatchPoint HRP Development Solution was added and incubated for 10 min in the dark while gently shaking. The fluorescence was measured using BioTek Synergy H1 plate reader at excitation at 530 nm and emission at 590 nm.

### Immunoblot

Platelets were seeded on 8-well chamber cover glass slides coated with Collagen I. After exposing the platelets with 20 μg/ml spike protein or its vehicle control, the floating and adherent platelets were separately lysed in NP-40 lysis buffer (50 mM Tris-HCl pH 7.4, 150 mM NaCl, 1 mM NaF, 1 mM Na3VO4, 1% IGEPAL^®^ CA-630, 1 mM EDTA, protease inhibitors). The total protein concentration was determined using the Pierce™ Rapid Gold BCA Protein Assay Kit (Thermo Fisher #53226). 30 μg of total protein sample was mixed with Laemmli buffer (180 mM DTT (Sigma #D9779), 4% SDS (VWR #442444H), 160 mM Tris-HCl pH 6.8, 20% glycerol (VWR #24388.295), bromophenol blue). The samples were then heated at 95°C for 5 min, spun down and were loaded on a BoltTM 4-12% Bis-tris (Invitrogen #NW04120BOX), and SDS-PAGE was run at 180 V for 40 min in 1X MES buffer and subsequently transferred to a PVDF membrane using the Bio-Rad Trans-Blot^®^ Turbo system according to the manufacturer’s instructions. The membrane was incubated with 4% ECL Blocking Agent (Fisher Scientific # 45001197) prepared in 1x TBST buffer (20 mM Tris-HCl, 150 mM NaCl, 0.1% Tween^®^ 20) at 4°C for 3 h. When blotting against phospho-antibodies, the membrane was incubated with 4% BLOCK ACE (Bio-Rad # BUF029). The primary antibodies (Phospho-FAK (Tyr397) Polyclonal Antibody (Thermo Fisher, 44-624G; 1:1000), Anti-GAPDH antibody Mouse monoclonal (Sigma #8795; 1:1000) for control were incubated at 4°C overnight. After three washes with TBST, the membranes were incubated with secondary horseradish peroxidase-coupled goat anti-rabbit or anti-mouse antibody (1:10,000; Bethyl) for 60 min. The membranes were washed in TBST and the chemiluminescence signal was revealed by incubating with ECL substrate (Bio-Rad) for 2 min and imaged using Amersham™ Imager 600. The signals were quantified using FIJI and they were normalized using GAPDH signals. Anti-GAPDH antibody was used as a control as regulation of cytoskeleton components such as actin and tubulin may occur upon activation of platelet.

### Pseudotyped SARS-CoV-2 S lentiviral particle production and transduction

Pseudotyped S lentivirus particles were either purchased (BPI #79981-1), obtained from BEI resources (BEI #NR-53818), NIAID, NIH or produced by adapting the published protocols^51,57^. Necessary reagents for the production were obtained from BEI resources, NIAID, NIH. Briefly, X-Lenti 293T cells (Takara #632180) plated in 6-well plates were co-transfected with 1 μg of lentiviral backbone ZsGreen (BEI #NR-52516); 0.22 μg each of the helper plasmids HDM-Hgpm2 (BEI #NR-52517), pRC-CMV-Rev1b (BEI #NR-52519), HDM-tat1b (BEI #NR-52518); and 0.34 μg of viral entry protein (SARS-CoV-2 Spike, BEI #NR-53742) or pCMV-VSVG (a gift from B. Weinberg, Addgene plasmid #8454) using 8 μl of TransIT-293 (Mirus Bio #MIR 2705) transfection reagent per well. On the next day, media was exchanged and collected after 60 h post transfection. The virus-containing supernatants were passed through a 0.45 μm filter and stored at −80°C. To test the transducing ability of the viral particles, different volumes of the viral supernatant were added to HEK-293T-hACE2 (BEI #NR-52511, HEK-293T cells constitutively expressing ACE2) for 60 h and were analyzed for GFP-positive cells using a BD LSRFortessa Cell Analyzer. For negative stain experiments, virus was produced in large quantities, concentrated using ultracentrifugation on a 20% sucrose layer^58^.

### Negative Staining of Pseudotyped SARS-CoV-2 S lentiviral particle

10 μl of concentrated Pseudotyped SARS-CoV-2 S lentiviral particles were incubated for 30 min on carbon coated grids (EMS #CF200-CU). Excess solution was removed by back blotting, the sample was washed three times in a drop of 1xPBS and H_2_O, subsequently stained with 5 μl of 2% (w/v) uranyl acetate solution. The grids were imaged using a Tecnai T12 transmission electron microscope (FEI) operated at 120 keV.

### Cryo-electron tomography sample preparation and data acquisition

Quantifoil grids (MultiA Au200 & SiO_2_ R1/4 Au200) were glow discharged using a Pelco easiGlow™ at negative discharge, 15 mA plasma current and 0.38 mbar residual air pressure for 45 s. The glow discharged grids were placed in a cell culture dish (greiner #627170), covered in diluted coating solution, and incubated. After coating, the grids were washed in PBS and blocked with 1% BSA in PBS. After blocking, grids were covered in Tyrode’s buffer. Platelets, preincubated with 9.75 μg/ml S protein or control buffer (20 mM HEPES, 150 mM NaCl, 0.01% LMNG, pH 7.5) for 4 h, were added to cover the grids using a wide orifice and adhered for 1 h. 3 μl Tyrode’s buffer was added to grids with adhered platelets and vitrified in liquid ethane using a Thermo Fisher Scientific Vitrobot MarkVI, conditioned at 37 °C and 100% humidity.

The data was collected using a Titan Krios (Thermo Fischer Scientific), equipped with a Gatan Quantum 967 LS and K3 Summit direct detector at an acceleration voltage of 300kV. Tilt-series were collected from −60° to 60° with 2° angular increment with a defocus range between −3 μm to −5 μm using a dose-symmetric acquisition scheme in SERIAL-EM software^59^. At a nominal magnification of 33,000 x, corresponding to a final pixel size of 2.76 Å. The total accumulated electron dose was 123 e^-^/Å^2^. Images were acquired as six-frame movies in super-resolution mode. A total of 8 tilt-series were assessed for this study.

### Cryo-electron tomography reconstruction and segmentation

Images were motion-corrected and filtered according to their cumulative dose using the software MotionCor2^60^. The tilt-series was aligned using the IMOD ETOMO package^61^. Tomograms were reconstructed, unbinned and 4x binned, from aligned stacks as weighted back-projection in IMOD. The contrast of the tomograms was increased by applying Matlab based deconvolution filter^62^. The CTF was estimated using GCTF^63^ and then the IMOD ctfphaseflip implementation was used for phase correction^64^.

Tomograms were manually segmented using AMIRA (Thermo Fisher Scientific) or 3DMOD (IMOD). The data was plotted using GraphPad PRISM.

### Actin analysis

For the actin analysis, we analyzed 282 actin filaments. Actin was segmented manually using IMOD software^61^. The IMOD model files were converted into coordinate files using IMOD (model2point). Each actin filament was segmented into 3nm and 10nm spaced segments for further analysis. For each actin filament the following two parameters were calculated: a) angle of individual 10 nm actin segment against the longitudinal axis of platelets protrusion and b) length of each actin filament. For determination of angle, a vector pointing towards the longitudinal axis of platelets is taken as reference vector and angle between this vector and each actin segment was calculated using Python3 NumPy library. Similarly, the length of each actin filament was calculated by adding the distance of all the segment present in each filament by using Python NumPy and SciPy libraries. The data was shown as histogram plots using GraphPad PRISM.

### Subtomogram averaging of S protein and distance analysis

4167 S protein particles were manually picked from 4-times binned tomograms using IMOD softwared (3dmod). The coordinates of the picked particles were then transferred to RELION-3^65^, and particles were extracted to a box with the size of 120 pixel from the original, unbinned electron tomograms. The extracted particles are located on the membrane surface and the membrane densities can interfere with the alignment process. To computationally supress the membrane density, Pyseg^49^ scheme was used. Pyseg uses a basis of discrete Morse theory ^66,67^. Initially, all subvolumes where aligned with respect to the membrane using RELION3 as its density is stronger than the density of the protein. Afterwards, membrane densities for each subvolume are suppressed by assigning random background values to membrane voxels, membrane and background voxels are identified with an input mask. The initial template used for the alignment was our SPA 3D reconstruction of S protein but low pass filtered to 60 Å. At this resolution, only general shape and size of S protein was visible. The initial alignment was done using 3D auto-refine and 3D classification schemes from RELION-3. The particles were divided into 4 classes by 3D classification and the classes that showed the remaining membrane densities and noises were discarded for the further reconstruction. The class that showed the most features was selected and the final reconstruction was performed using 976 particles with C3 symmetry and with the mask that is created from the reconstruction of the previous run. The resolution was estimated by comparing the FSC of two separately computed averages from odd and even half-sets from the final refinement, the standard procedure available from RELION-3. The final resolution was estimated to be 13.8 Å with the FSC 0.143 criterion. The refinement without C3 symmetry was also performed, revealing the resolution of 20 Å.

Distances between S protein particles were calculated from the coordinates using Python3 (numpy and scipy libraries)^68^. Positions of S protein were defined by manual picking. For each particle, the closest neighboring distance was plotted into the distance distribution histogram using GraphPad PRISM.

### Membrane Curvature analysis and determination of S protein orientation and distance from the membrane

The segmentation of the membrane was done manually using AMIRA (Thermo Fisher) software. The membrane curvature was then determined using python based software PyCurv^69^ using the standard workflow (https://github.com/kalemaria/pycurv). The segmented membrane from the binned tomogram was used as the input. The software then converts this segmentation, i.e., a set of voxels, into a surface, mesh of triangles. This surface of triangular mesh is converted into a surface graph, normal vectors and local curvature was then computed for every triangle center.

In order to measure the distance between the density corresponding to S protein and the membrane, the Euclidean distance between the refined coordinates of S protein particle to all the triangles on the membrane surface was calculated and the smallest distance was considered. Similarly, to determine the orientation of S protein with respect to the membrane, the angle between the vector pointing towards the longest axis of S protein and the normal vector of the closest triangle was calculated (Fig. 4D-E). Both distance and angular orientation was calculated using Python3 (numpy and scipy libraries). The position and orientation of S protein was visually assessed by the Place Object plug-in^70^ in Chimera^71^.

### Single particle analysis of SARS-Cov-2 S protein, data collection and image analysis

SARS-CoV-2 S protein (Cube Biotech) was recorded at 0.3 mg/ml. 3-μl of sample was applied on glow discharged (Pelco easiGlow™; t=20 s; I=20 mA) 200-mesh R1.2/1.3 Quantifoil girds. After blotting for 3.5 s at 4°C and 100% humidity, the sample was vitrified in liquid ethane using a Vitrobot Mark IV.

The data was acquired on a Glacios (Thermo Fisher Scientific) operated at 200 keV and equipped with a Falcon 4 direct electron detector. Images were collected by EPU software (Thermo Fisher Scientific) with a pixel size of 0.93 Å with a defocus range from −0.8 to −2.4 μm. In total, 3060 movies, divided in 40 frames, were collected with a total dose of 49 e^-^/Å^2^.

All data processing steps were performed in cryoSAPRC v.3.3.1^72^. Motion correction in patch mode, CTF estimation in patch mode and subsequent blob picking were performed. An initial set of obtained particles was used for training a Topaz which was optimized over several rounds to extract the final set of 11,549 particles^73^. Heterogenous refinement was performed to separate open and closed states of S proteins. The closed-state S protein class was further refined using C3 symmetry. The final map was reconstructed by non-uniform refinement with per particle CTF estimation and aberration correction from 7,618 particles^74,75^ at a resolution of 3.56 Å with the FSC 0.143 criterion. The final map was sharpened using DeepEMhancer^76^ and visualized in ChimeraX^77^. For the reconstruction of S protein in the open conformation, the same analysis scheme was applied using C1 symmetry on a final set of 3,931 particles. The final resolution was estimated to be 7.44 Å with the FSC 0.143 criterion.

### Ligand Binding ELISA Assay

Ligand binding assays were performed as described previously^78^. Solutions of human plasma vitronectin (5 μg/ml), human fibrinogen (20 μg/ml), bovine plasma fibronectin (10 μg/ml), S protein (residues 1-1212 with stabilizing mutations R684G, R685S, R678G, K998P, and V999P) with a C-terminal foldon motif and a His6 tag, 20 μg/ml)^79^ and SARS-CoV-2 S1(residues 1-696 with a C-terminal foldon motif and a His6 tag, 20 μg/ml) in TBS were used to coat 96-well polyvinylchloride microtiter plates (Nunc Maxisorp #44-2404-21) for 6 h at RT. Non-coated well was used to determine unspecific binding background values. After blocking overnight at 4°C (Blocking One, Nacalai #03953-95), velcro-tagged integrins α_v_β_3_, α_IIb_β_3_, and α_5_β_1_ diluted at 10 μg/ml in 20 mM Hepes, 150 mM NaCl, 1 mM MnCl_2_, pH 7.2 were incubated for 2 h at RT to allow binding to the immobilized ligands. Bound integrins were quantified by an enzyme-linked immunosorbent (ELISA)-like solid-phase assay using biotinylated rabbit anti-velcro (against ACIS/BASE coiled-coil) antibody and HRP-conjugated streptavidin (VECTOR Laboratories #SA-5004). After addition of ABTS, readout was performed at 405 nm.

## Figure Legends

**Figure S1. Platelets incubated with SARS-CoV-2 S protein reveal proplatelet-like morphologies.** (A) A subset of platelets showed a tubular appearance with multiple globular bodies along their elongated shape after the incubation with S protein. Some platelets had a ring-shaped appearance in the presence of the S protein. (B) The adherent platelets were collected in RIPA lysis buffer and the total concentration was quantified with Bradford assay. 30 μg of total protein was loaded on the gel and probed against pFAK or GAPDH. (C) Quantification of pFAK in adherent platelets normalized against GAPDH concentrations, showing the values of 2.04 (donor 1), 1.22 (donor 2) and 1.09 (donor 3). (D) The floating platelets were handled as specified in (B) and probed against pFAK, MLC or GAPDH. Scale bar: 5 μm.

**Figure S2. Cryo-electron tomographic visualization of platelets incubated exposed to SARS-CoV-2 S protein.** (A) Filopodia width of the platelet incubated with S protein. (B) Central Slices of analyzed tomograms. The images show a slice through the deconvoluted tomogram used for further analysis. Scale Bars: (A) = 100 nm; (B) = 200 nm

**Figure S3. Cryo-EM structure of the SARS-CoV-2 S protein.** (A) Cryo-EM map of S protein in the open conformation at a resolution of 7.44 Å. (B) Cryo-EM map of S protein in the closed conformation at a resolution of 3.56Å at FSC=0.143. (C) Gold standard FSC curves of the reconstructed S protein in the open and closed conformation. (D) (E) Gold standard FSC curve of the sub-tomogram averaged S protein reconstruction in the closed conformation with an estimated resolution of 13.8 Å at FSC=0.143.

**Figure S4. Characterization of SARS-CoV-2 S-pseudotyped lentiviral particles.** (A) Negative-staining EM with 2% uranyl acetate of SARS-CoV-2 S-pseudotyped lentiviral particles. (B) Light microscopic images of HEK-hACE2 cells with or without treatment with pseudotyped lentivirus encoding ZsGreen. (C) Flowcytometry analysis of HEK-hACE2 cells transduced with pseudotyped lentivirus encoding ZsGreen backbone plasmid. The plot shows percentage of green-fluorescent cells with or without the incubation of S-pseudotyped lentivirus with HEK-hACE2 cells. The gate was set that the uninfected cells show less than 1% positive cells and the same gate has been applied to infected cells. Scale bars: (A) = 100 nm; (B) = 50 μm.

**Figure S5. SARS-CoV-2 S-pseudotyped viral particles on platelet plasma membrane by cryo-ET.** (A) Extracellular vesicles and S protein pseudotyped virus under cryo-EM condition. (E: extracellular, I: intracellular). (B) Overview of grid square of platelets incubated with pseudotyped viral particles. (C) Low magnification of platelets observed in the presence of pseudotyped viral particles. (D) Platelet with pseudotyped viral particle exhibits the formation of a filopodial protrusion. The dashed box indicates the area of tomogram data collection. (E) Slice of the reconstructed tomogram showing a virus-like particle on the platelet plasma membrane. (F) Segmentation of the tomogram with virus-like particle at the platelet plasma membrane (purple – platelet plasma membrane, yellow – virus-like particle membrane). The tomogram lacks top and bottom due to the “missing wedge” effect of tomographic data collection. (G) Zoom-in views on the contact site of platelet and virus-like particle. (H) Zoom-in view on the segmented platelet membrane and virus-like particle membrane (purple – platelet plasma membrane, yellow – virus-like particle membrane) (I) Slices thorough the virus-like particle at the filopodia tip. Red arrows point at the protein densities on the membrane surface. Scale bars: (A)=100nm; (B)=10 μm; (C)= 5 μm; (D)=1 μm; (E)&(F)=200 nm; (G)&(I)=20nm.

**Movie S1. Representative DIC time lapse movies of platelets with or without SARS-CoV-2 S protein.** (A) Movies of platelets seeded on collagen type I coating preincubated without (left) or with S protein (right). (B) Movies of platelets seeded on poly-lysine coating preincubated without (left) or with S protein (right). (C) Movies of platelets seeded on fibronectin coating preincubated without (left) or with S protein (right). Scale bars: 5 μm.

**Movie S2. Reconstructed tomograms, acquired on platelets incubated with SARS-CoV-2 S protein.** (A) Tomographic reconstruction of a filopodial platelet protrusion. (B) Tomographic reconstruction of a platelet protrusion including a microtubule. Scale Bars: (A) 100 nm; (B) 200 nm.

