## Supplementary figures and images for "Direct Cryo-ET observation of platelet deformation induced by SARS-CoV-2 Spike protein"

### Supplementary Material

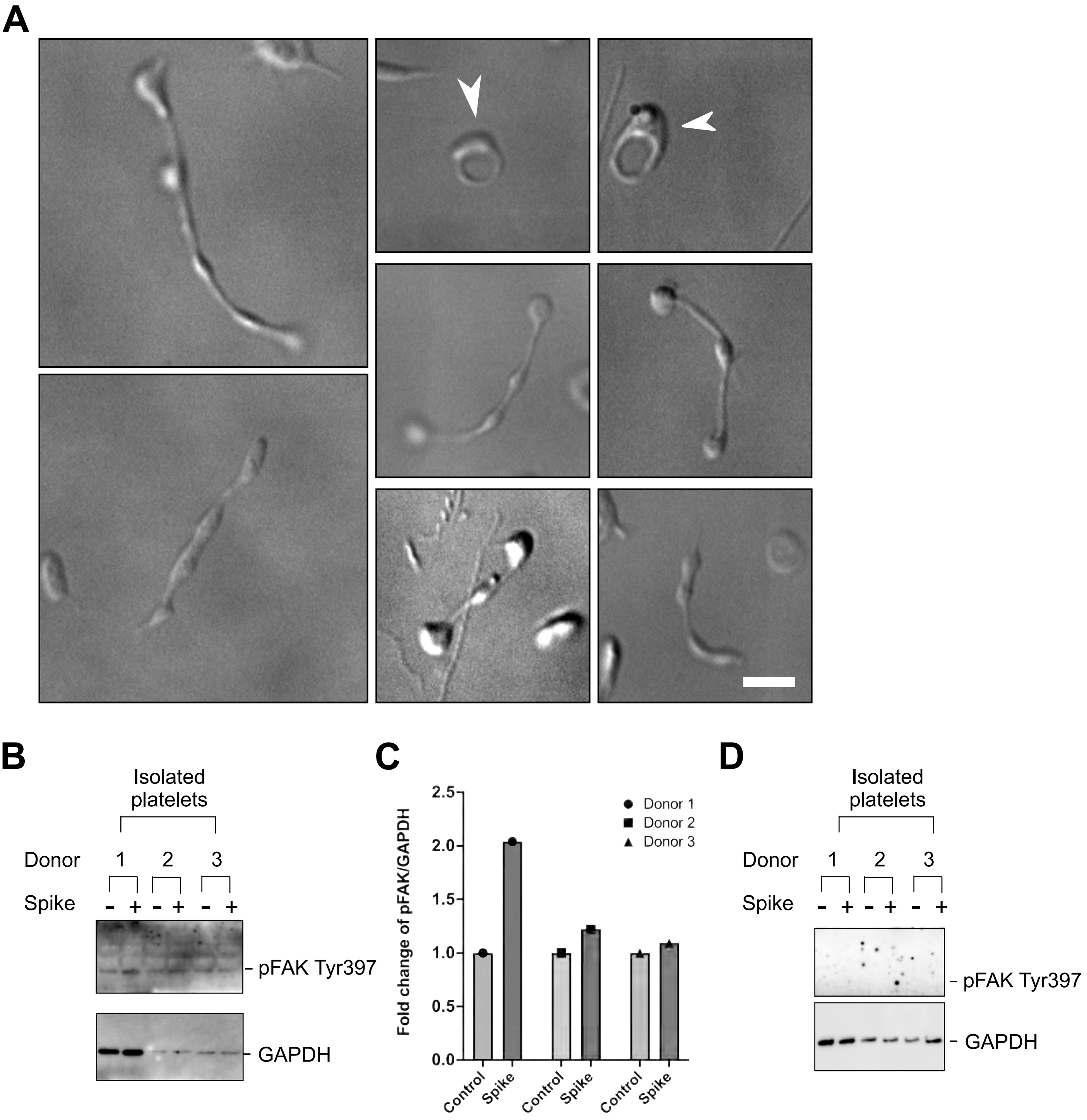

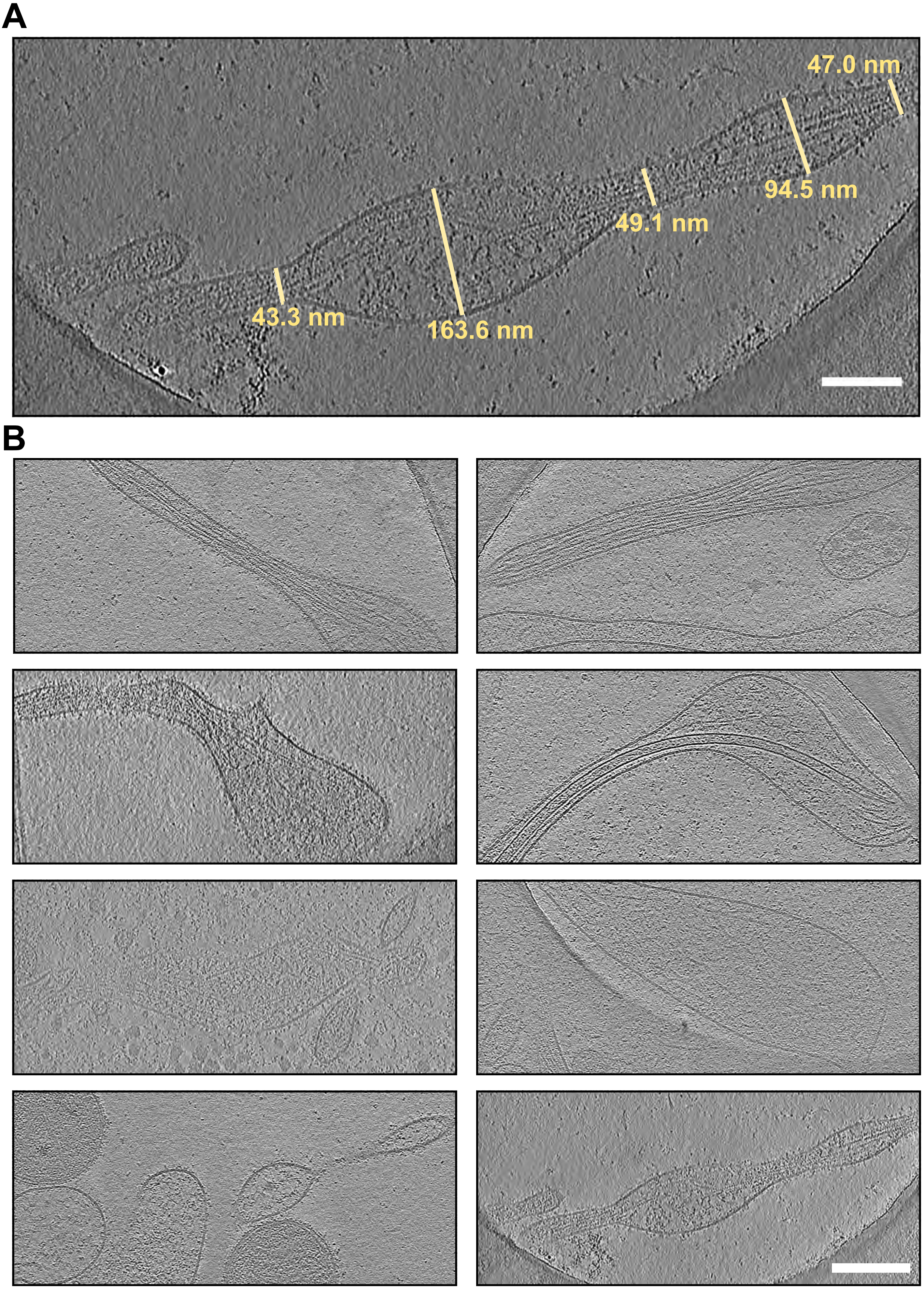

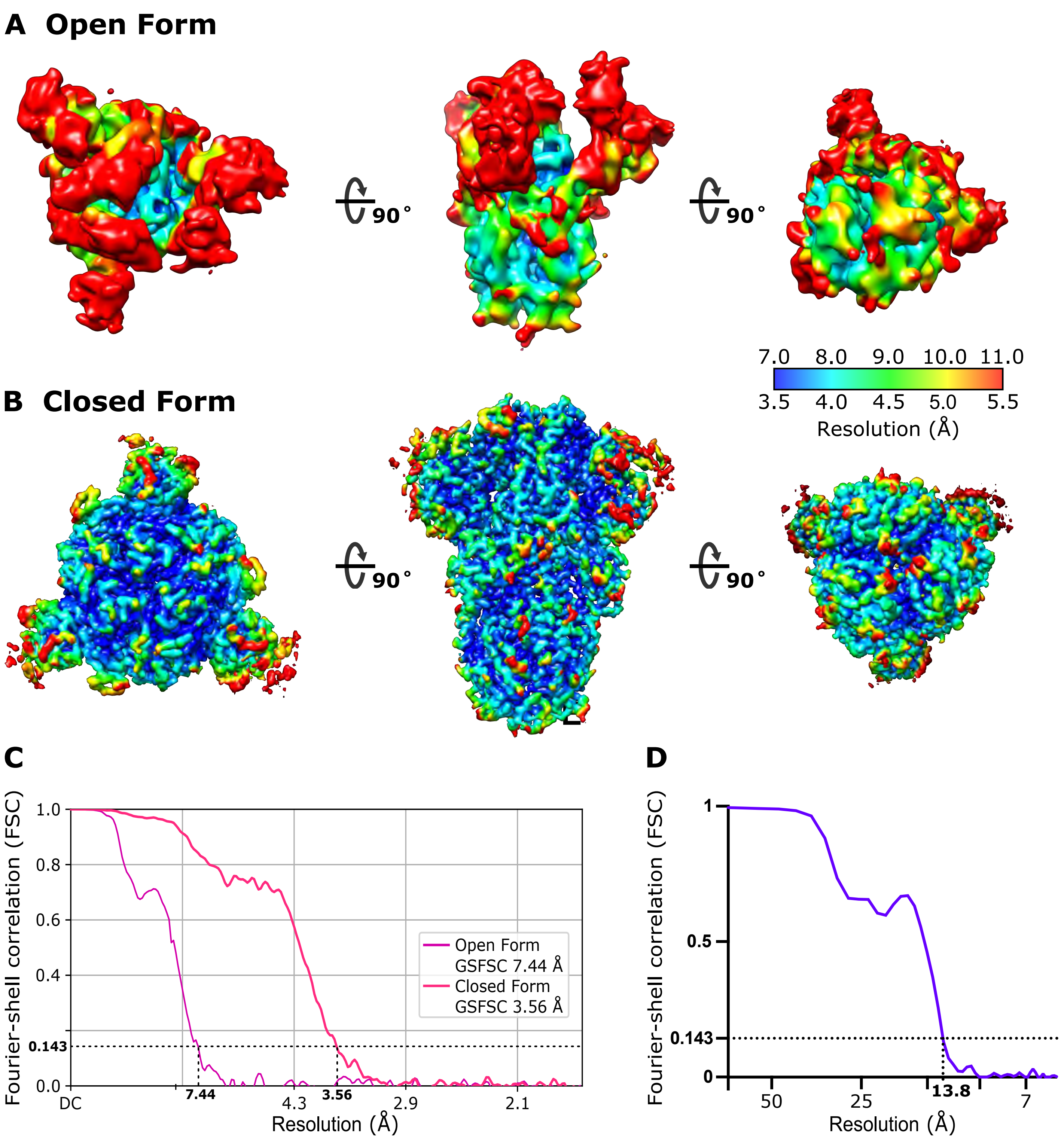

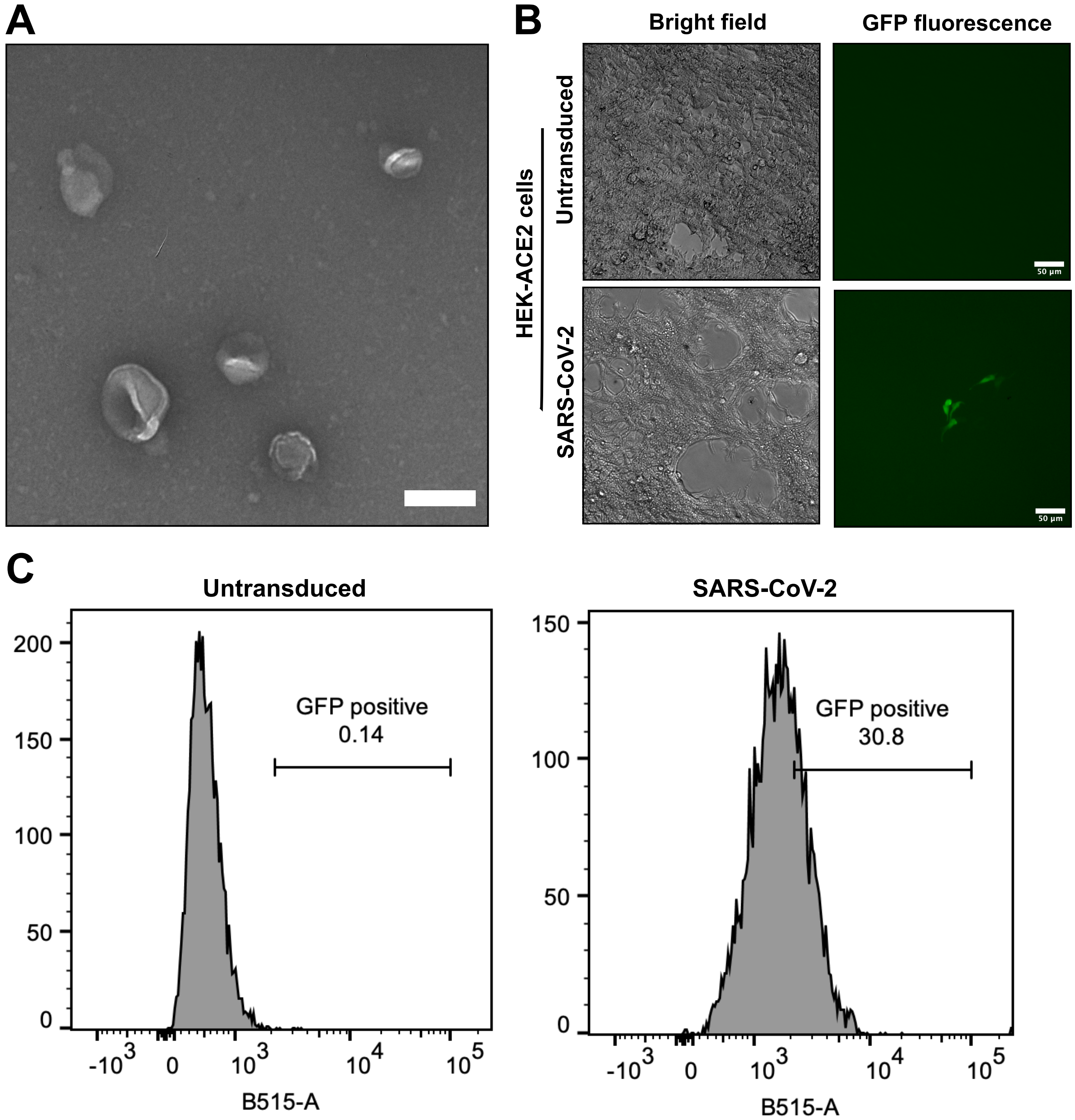

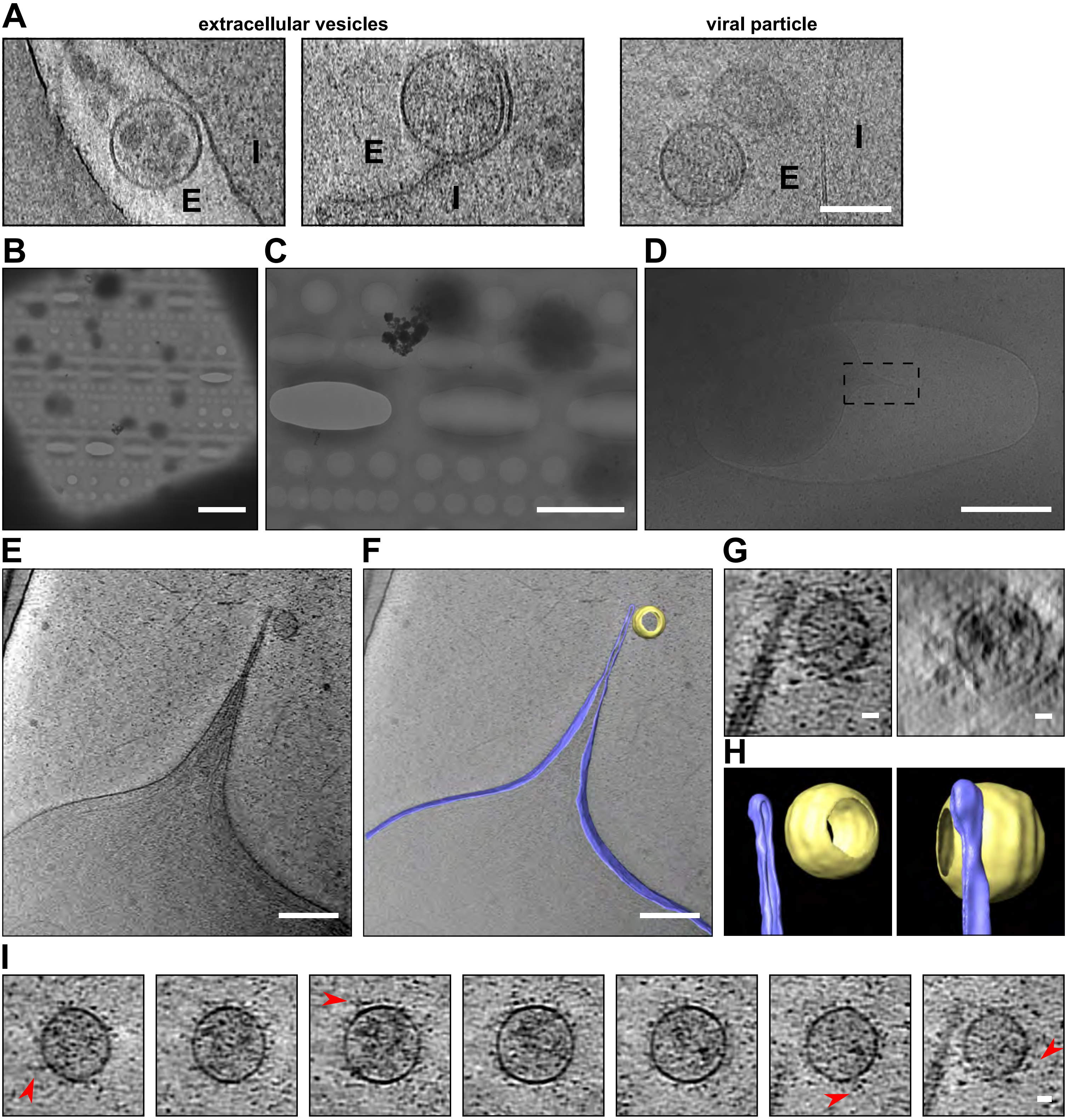
